# SPOT-RASTR - a cryo-EM specimen preparation technique that overcomes problems with preferred orientation and the air/water interface

**DOI:** 10.1101/2024.01.24.577038

**Authors:** Behrouz G. Esfahani, Peter S. Randolph, Ruizhi Peng, Timothy Grant, M. Elizabeth Stroupe, Scott M. Stagg

## Abstract

In cryogenic electron microscopy (cryo-EM), specimen preparation remains a bottleneck despite recent advancements. Classical plunge freezing methods often result in issues like aggregation and preferred orientations at the air/water interface. Many alternative methods have been proposed, but there remains a lack a universal solution, and multiple techniques are often required for challenging samples. Here, we demonstrate the use of lipid nanotubes with nickel NTA headgroups as a platform for cryo-EM sample preparation. His-tagged specimens of interest are added to the tubules, and they can be frozen by conventional plunge freezing. We show that the nanotubes protect samples from the air/water interface and promote a wider range of orientations. The reconstruction of average subtracted tubular regions (RASTR) method allows for the removal of the nanotubule signal from the cryo-EM images resulting in isolated images of specimens of interest. Testing with β-galactosidase validates the method’s ability to capture particles at lower concentrations, overcome preferred orientations, and achieve near-atomic resolution reconstructions. Since the nanotubules can be identified and targeted automatically at low magnification, the method enables fully automated data collection. Furthermore, the particles on the tubes can be automatically identified and centered using 2D classification enabling particle picking without requiring prior information. Altogether, our approach that we call specimen preparation on a tube RASTR (SPOT-RASTR) holds promise for overcoming air-water interface and preferred orientation challenges and offers the potential for fully automated cryo-EM data collection and structure determination.

## Introduction

Recent technological innovations have significantly increased the efficiency of cryo-EM, causing a proliferation in the number of structures resolved with ever-increasing resolution. As many of the “low hanging fruit” structures have been determined, studies have shifted toward increasingly complex specimens, pushing the capabilities of current processing software, and requiring novel methodologies. Sample preparation in cryo-EM is often fraught – many samples require very high concentrations, do not go into holes in the EM substrate, and/or are degraded by interaction with the air/water interface (Noble, Wei, et al. 2018). Typically, samples are prepared for cryo-EM by placing them on a holey grid, blotting to a thin layer ^∼^100 nm thick or less, and freezing by plunging into liquid ethane. During the time that the sample is blotting (typically around 3 seconds), the sample can interact with the air/water interface 3000 times or more (Taylor and Glaeser 2008). These interactions with the air/water interface often introduce artifacts such as preferred orientations or in the worst cases entirely denature the sample. Recent studies have shown that in the majority of cryo-EM grids, the sample is located at the air water interface(Noble, Dandey, et al. 2018). Complicating the situation, every different sample whether its protein, nucleoprotein or large virus-like particle may behave differently while plunge-freezing (B.-G. Han et al. 2023; B. G. Han et al. 2022; Weissenberger, Henderikx, and Peters 2021). These problems including aggregation, denaturation, disassembly, preferred orientation, etc. are mostly caused by the air-water interface binding of the sample during freezing (Chen et al. 2019) (Fig.S1).

Many strategies have been developed to improve the quality of the final sample in cryo-EM, and sample preparation can be optimized at every stage of the vitrification workflow. For instance, sample preparation can be optimized at the biochemical level to have as uniform and homogenous sample as possible (Glaeser 2021; Stark 2010), the grid selection level to have bigger or smaller hole size, or by selecting a different support material such as gold or carbon (Park et al. 2020). Other sample preservation approaches have been used such as glutaraldehyde fixation or fixation in agar which may help to stabilize heterogenous samples (Adamus et al. 2019). Many samples have seen success by using a binding surface like graphene or graphene oxide to trap the sample and keep it away from the air-water interface (Chen et al. 2019; D’Imprima et al. 2019). Alternatively some approaches have used fast automated plunging with self-wicking grids to have a much faster plunge/blot time and involve less human error during the freezing (Levitz et al. 2022). Each of these approaches address different challenges in the preparation pathway, but a universal solution has not yet been found, leaving space for the development of new tools.

Tubular samples offer an attractive platform for overcoming the challenges of cryo-EM specimen preparation. A tubule decorated with a protein of interest, would: 1) display particles in all required orientations, and 2) hold the protein away from the damaging air/water interface. Although a small percentage of particles may still contact the air/water interface, the structural information from these views will be provided from the decorated particles on the other side of the tube. Kubalek and colleagues have shown that the lipid galactosyl ceramide (GalCer) (Fig. 1A) when added to phospholipids will tubulate them into multi-micron long tubules (Wilson-Kubalek et al. 1998). These can be doped with functionalized lipids such as those with Nickel-NTA headgroups (DGS-NTA) (Fig. 1A) that can bind His-tagged proteins (Raghunath and Dyer 2019). Kubalek et al. proposed that Ni^+^ doped GalCer tubes could be use as substrates for helical crystallization of His-tagged proteins and membrane proteins. Their method required proteins to crystalize on the tubular surface, and this technical challenge has limited adoption of the technique.

**Figure 1.**
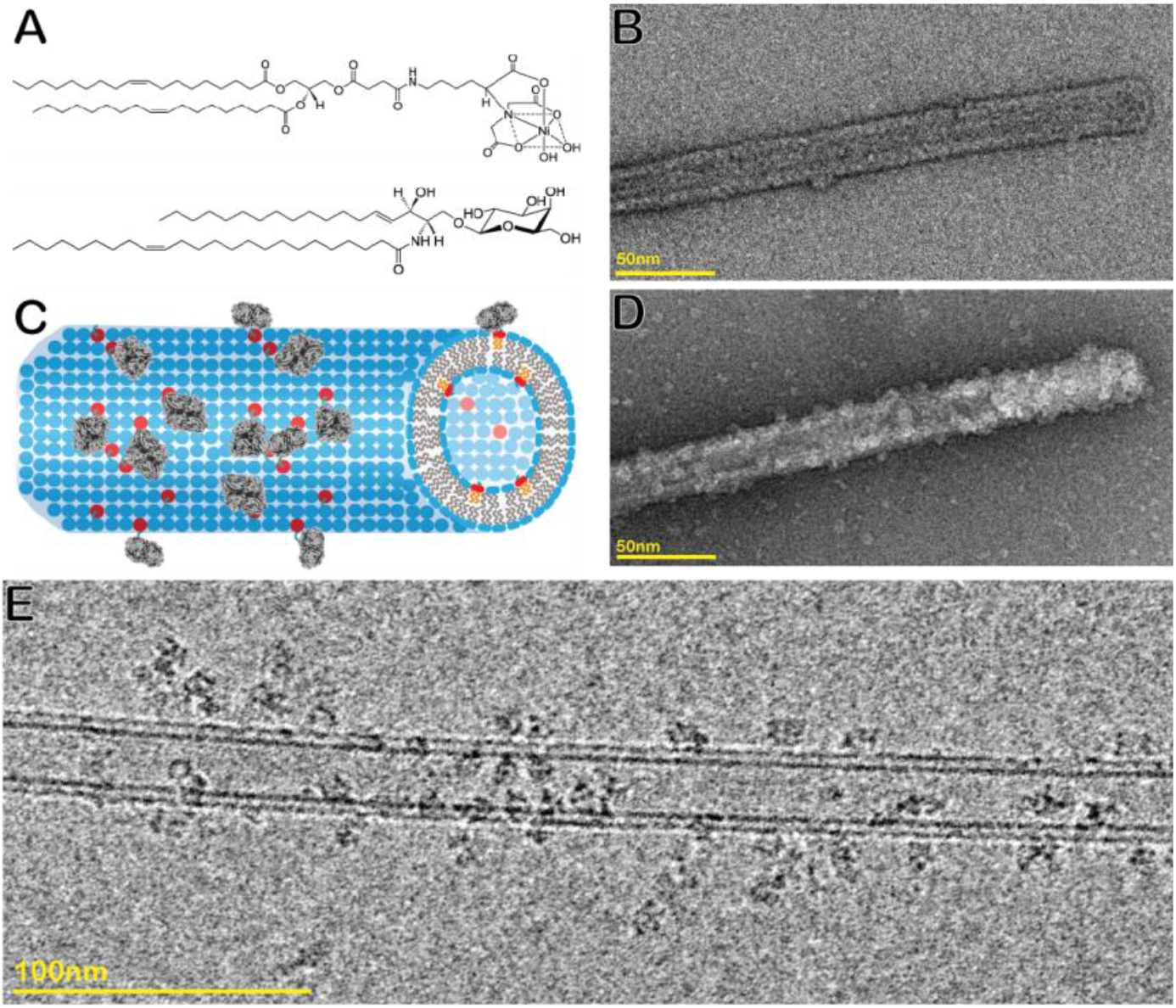
A) tube lipid content, top DGS-NTA lipids and bottom GalCer lipids. B) tubes negatively stained. C) Schematic tubes bound to β-gal. D) decorated tube with β-gal negatively stained. E) decorated tube with β-gal, cryoEM micrograph.

We recently published a method called reconstruction of average subtracted tubular regions (RASTR) that enables structure determination of challenging tubular samples (Randolph and Stagg 2020). In that method, we computationally align tubular samples and generate a so-called 3D azimuthal average that represents the rotational average density of tubular samples around their tubular axis. The 3D azimuthal average can be projected back in 2D and subtracted from the original 2D images to downweight or erase tubular features. Low complexity features such as membranes are subtracted particularly well using this approach. We have combined this approach together with Ni^+^ doped membrane nanotubes to create a new method called specimen preparation on a tube RASTR (SPOT-RASTR). In SPOT-RASTR, we sparsely bind His-tagged proteins of interest to the nanotubules (Fig. 1C) and use the RASTR process to computationally erase the nanotube membrane leaving images of the protein of interest. These can then be treated by conventional single particle methods to generate high-resolution reconstructions of the proteins. Here we demonstrated the efficacy of SPOT-RASTR using the 520 kDa protein β-galactosidase (β-gal). We show that our approach has many benefits including capturing and concentrating the sample, holding the sample away from the air/water interface, and filling in missing views compared to conventional cryo-EM preparation. We show that by picking the tubules alone, we can identify the β-gal particles in a completely automated user-free method that we call picking by classification. Finally, the tubules are easily identifiable at low magnification and can be automatically targeted. When combined, the SPOT-RASTR approach provides a workflow for completely automated structure determination from specimen preparation to 3D reconstruction.

## Results

### Nanotube preparation and optimization

The first step in our approach was to determine an optimal amount of Ni^+^ lipids to use in the tubule preparations. Tubule preparations with a 9 to 1 molar mixture of GalCer lipids to Ni^+^ lipids were prepared, and 0.15 mg/ml β-gal was added to them. The samples were then assessed by negative stain EM. In the absence of β-gal, the tubules had a smooth appearance with widths between 25 and 30 nm, similar to what had been reported previously in the literature (Fig. 1B). When β-gal was added, the specimen could be observed decorating the surface of the tubules (Fig. 1D).

The next step was to prepare the tubules for cryo-EM. We initially tried adding the tubules together with the His-tagged β-gal and then using conventional plunge freezing with a Vitrobot Mark IV (ThermoFisher). Using this approach, we noticed that there were very few individual tubules, and there were large numbers of huge tubule aggregates (Fig. S2). We hypothesized that the tubules were aggregating because of the oligomeric nature of the β-gal. There are 4 β-gal monomers in each β-gal particle, thus a single β-gal molecule can crosslink multiple tubules. To overcome this problem, we first applied the tubules to the Quantifoil grids, then β-gal was added to the pre-adhered tubules. This was incubated for 10 s and finally plunge frozen. This resulted in a large number of individual tubules that were decorated with β-gal (Fig. 1E). The tubules could be clearly visualized at lower magnification (4800 X) which enabled targeting of tubules for high-throughput data collection.

The number of tubules per hole was relatively low, so we attempted to increase the concentration by applying nanotubes followed by blotting and then another round of tube application before adding the sample (β-gal). This resulted in increased tubule concentration (Fig. S3A), but another form of tube-like objects was also observed. These were nearly the same diameter as the nanotubules, but they had a ghostly appearance with lower contrast without the characteristic double lines of the bilayer (Fig. S3B, 3C, 3D). They appear more like incomplete tubules. Our best estimation is that these are the remnants of tubules that were ripped apart during blotting. These “ghost tubes” are similar to what occurs in the technique of cellular “unroofing” which tears off a patch of plasma membrane from adhered cells (Usukura et al. 2016). Consistent with this idea, the ghost tubes were frequently decorated with β-gal (Fig. S3B, S3C). We have seen the ghost tubes in all preparations, but they are more abundant when we apply the nanotube to grids twice to increase the total number of the tubes. After two tube application followed by side-blotting, we have found that single sided blotting (accomplished by having a layer of Teflon instead of filter paper for one of the Vitrobot’s blotting pads) on the same side of the carbon support to which we apply protein sample will also increase the number of the nanotubes sticking to the grid. Another type of tubes was very rare, highly dense, and seem deformed with no β-gal particles decorated on their surface that we call “dark tubes” (Fig. S3G).

After optimization, we found that decorated tubules were maximized, and aberrant tubules minimized by generating tubules using 9:1 of Ni^+^ GalCer lipids to lipids (DGS-NTA) molar ratio and adhering the tubules first to the grid before applying the protein sample. The method is highly reproducible, and in more than 20 repeats of plunging experiments where we started with application of the tubes to the grid, we observed only small variation from one grid to the next. The method performs similarly across different grid types such Ultrafoils and Quantifoils with different mesh and hole size (ranging from 1.2 um to 3.5 um). In each of these cases, the dispersion of the tubes was reasonable to collect data on. We found that the tubules were easier to identify and target using grids with a carbon mesh because the tubes are identifiable on the support and show more overall contrast, so these are what we collected datasets on.

### Particles are held away from the air/water interface and decorate all sides of the nanotubes in SPOT-RASTR

We collected cryo tomograms in order to characterize how the β-gal particles were distributed on the tubules and in the vitreous ice. ^∼^60 tilt series were collected, and these were aligned and reconstructed using AreTomo (S. Zheng et al. 2022) (Video. S1). Inspection of the resulting tomograms revealed that the decorated tubules had a few common characteristics: the diameter was consistent, they tend to disperse in different z heights, and they spanned across the ice instead of sticking to a single side. Most of the tubules were straight, and the location of the β-gal particles were random over the tubes (Fig. 2). Some tubules were embedded in the middle of the ice and had β-gal particles decorating all sides (Fig. S3). Occasionally one tubule would have one end anchored to the edge of a hole edge and the rest of the tubule was preserved in the vitreous ice at a slight angle. Also, occasionally tubules were observed with both ends anchored on the hole edges but with forming a belly embedded inside the ice (Fig. S3A). Some tubules traversed the air/water interface on one side of the grid to the other side, but the β-gal particles were still held away from the air/water interface. In the thinnest ice, the top and bottom of the tubules were located at the air/water interface. In these cases, the β-gal particles were not located at the top or bottom but were localized on the sides of the tubules. However, in these cases, the β-gal did not take on a single orientation. Instead, it appeared that while the β-gal was tethered to the tubule by its C-terminal His-tag, it was still free to rotate around the His tagged peptide, and thus a full 180° set of views of β-gal were achieved even with the particles that were bound to the edges of the tubules. In some areas there were clusters of tubes, but inspection of the tomograms revealed that they occupy different z heights inside thicker ice but with well-preserved particles attached. A final observation is that the distance between the β-gal protein and the membrane tube varied, and that may depend on the orientation of the particle relative to the position of the His tag. Altogether, we estimate that more than 90 percent of the tubes were embedded in ice regardless of ice thickness and were decorated with β-gal.

**Figure 2.**
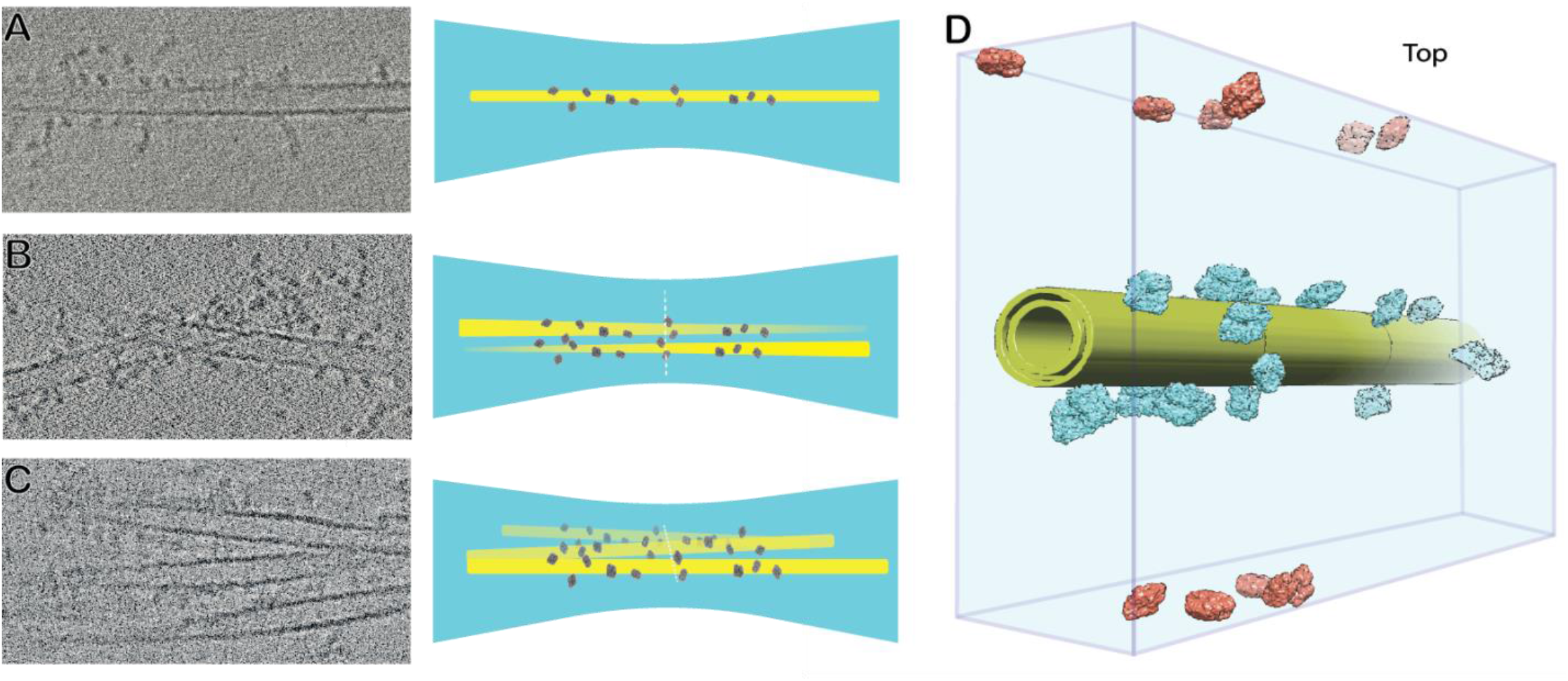
Frequent formation of the tubes on the cryo-EM grids identified by tomograms, A) single tube decorated with β-gal. B) two tubes crossing each other in different depths. C) tubes close together in the same plane. D) tomogram driven schematic from decorated tube, showing the free particles as well.

In addition to the particles bound to tubules, there were many free particles in the tomograms. These particles were mostly close or attached to the air/water interface. It is well-known that this can result in partial denaturation and/or preferred orientations. In our tomograms, it was difficult to determine whether they were denatured, but it was clear that certain views were preferred (Fig. 2D), and these were consistent with the preferred views for β-gal with conventional single particle cryo-EM.

### SPOT-RASTR single particle data collection and processing

Given that the β-gal particles bound to tubules were well-preserved in vitreous ice, we proceeded to collect single particle cryo-EM data. Approximately 4,000 motion-corrected movies were obtained of β-gal-decorated tubules (Table 1). The tubules were then picked using the filament picker in cryoSPARC. This resulted in around 700,000 overlapping tubular segments. This was followed by multiple rounds of 2D classification to eliminate around 470,000 low-quality segment class averages, resulting in 230,000 well-aligning tubular segments (Fig. 3A). The next step was to subtract the signal for the tubular membrane. This was achieved using the RASTR approach (Randolph and Stagg 2020). Briefly, tubules were aligned in 2D along the y-axis. This determined the on-plane rotation Euler angles required for a 3D reconstruction. The final Euler angle was randomized for the tubular segments and were reconstructed resulting in an “azimuthal average” that represents the average of the tubular density around the tubular axis (Fig. 3A, B). This served as a reference for another round of tubular alignment and azimuthal averaging. After this process converged, the azimuthal average was projected back in 2D and subtracted from the original particle images. This resulted in nearly complete erasure of the membrane tubule in most cases (Fig. 3C, D).

**Table 1.** Data collection parameters.

| Dataset | Scope | Camera | Total Dose (e-/Å) | Magnification | Pixel Size (Å) | No. of Movies | No. of tube picks | No. of protein particles | Symmetry Applied | Final Resolution (Å) |
| --- | --- | --- | --- | --- | --- | --- | --- | --- | --- | --- |
| β-gal on tubules | Titan Krios | GATAN K3 | 60 | 64,000 | 1.44 | 4,736 | 230,000 | 145,000 | D2 | 3.58 |
| SIRFP/HP on tubules | Titan Krios | Apollo | 60 | 59,000 | 0.765 | 5,847 | 48,000 | 70,000 | NA | NA |
| β-gal | Titan Krios | Apollo | 60 | 59,000 | 0.765 | 5,600 | NA | 210,000 | D2 | 2.03 |

**Figure 3.**
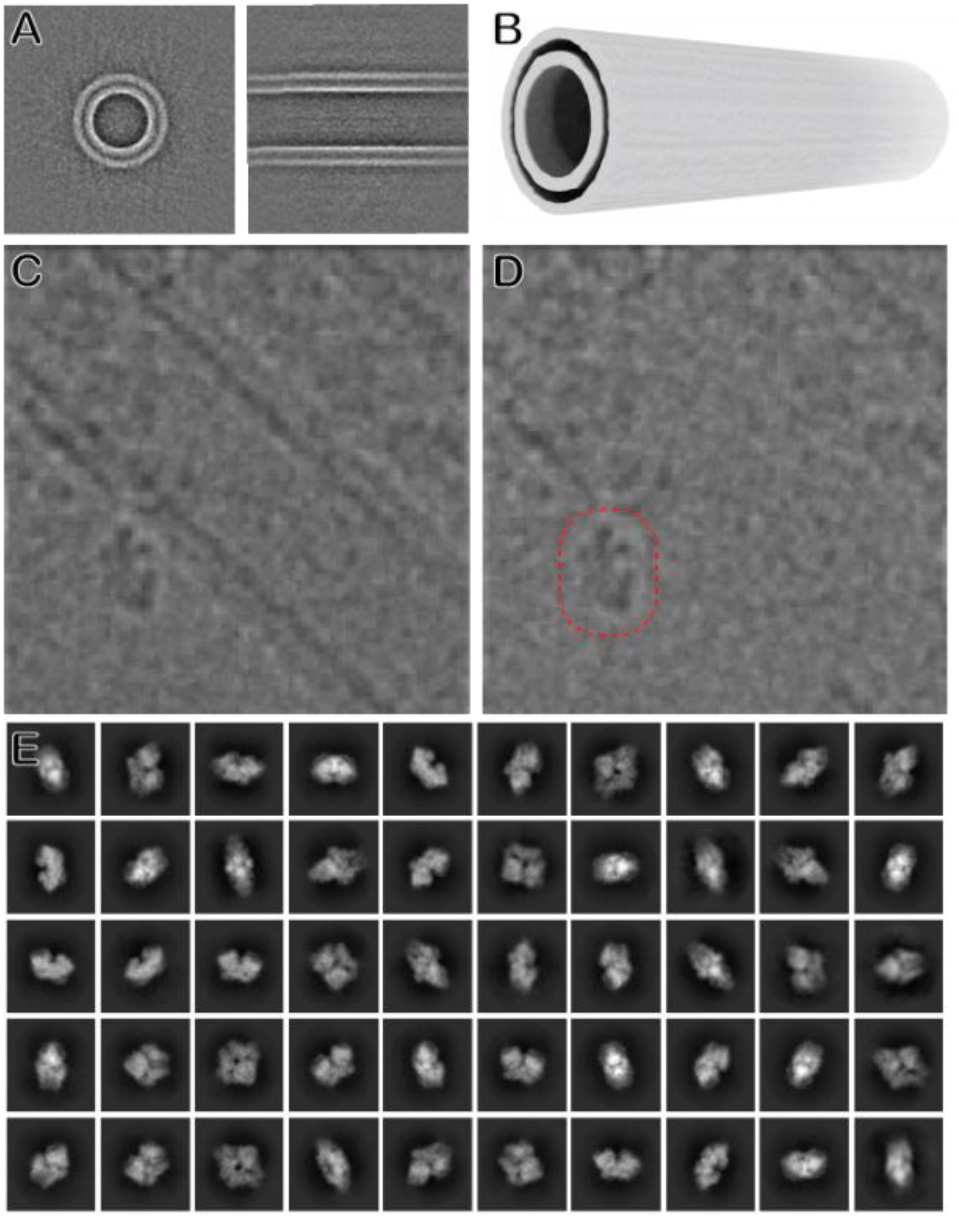
A) Different views of the aligned tube. B) 3D volume of the aligned tube to be subtracted from particles. C) tube pick showing the tube residue un-subtracted. D) same particle in C subtracted which is subtracted with B and picked particles using β-gal templates. E) β-gal 2D classes obtained from picked particles using the templates.

After the tubule signal subtraction, the next step was to box out the β-gal particles from the tubule-erased stack. The β-gal particles were picked using conventional template-based particle picking where each of the subtracted particles was considered as an individual micrograph and β-gal particles were identified by correlation against a β-gal template. This resulted in a new stack of centered β-gal particles. Since the original tubules were boxed out in overlapping segments, multiple identical β-gal particles were present in the β-gal stack. Thus, the duplicates were identified and eliminated using cross correlation. This resulted in a stack of ^∼^150,000 unique β-gal particles.

### SPOT-RASTR fills in missing Euler angles

The SPOT-RASTR particles were subjected to 3D single particle analysis in cryoSPARC, resulting in a 3.5 Å resolution map (Fig. 4A). PDB model 3j7h was refined into the map using PHENIX (Liebschner et al. 2019) (Table.2). No significant change was observed in the model comparing to the other models of the β-gal (see below). No evidence of the lipid trace on the map was observed indicating that the tubule signal was successfully subtracted. There were some particles where the tubule was not completely subtracted, but these were removed during the final 2D classification stage before the 3D reconstruction.

**Table 2.**
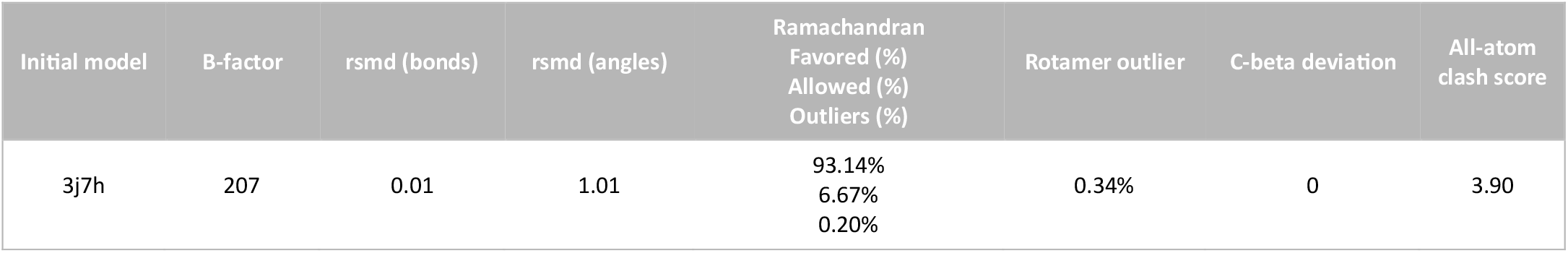
PHENIX refinement outputs.

| Initial model | B-factor | rmsd (bonds) | rmsd (angles) | Ramachandran Favored (%)<br>Allowed (%)<br>Outliers (%) | Rotamer outlier | C-beta deviation | All-atom clash score |
| --- | --- | --- | --- | --- | --- | --- | --- |
| 3j7h | 207 | 0.01 | 1.01 | 93.14%<br>6.67%<br>0.20% | 0.34% | 0 | 3.90 |

**Figure 4.**
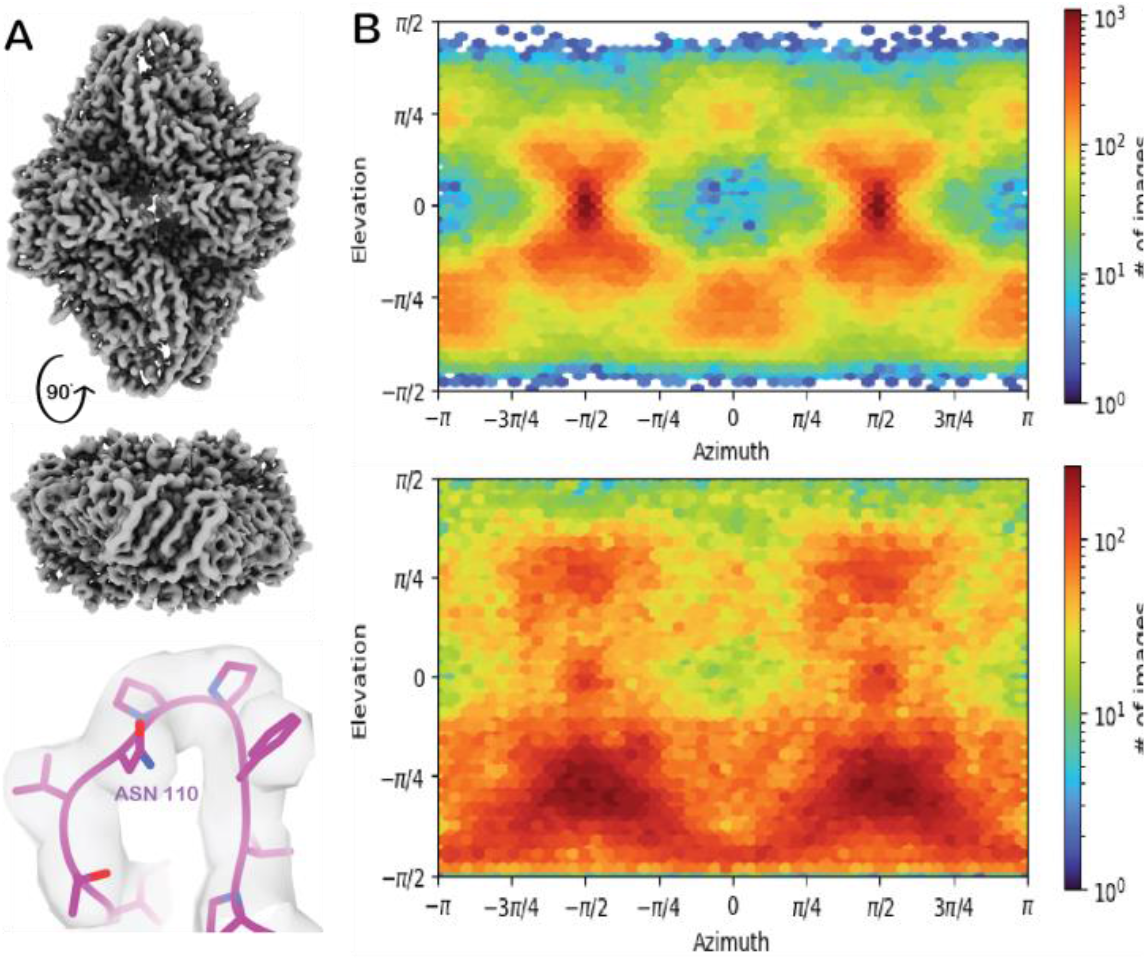
A) β-gal 3.5 Å resolution map and the fit quality. B) Particle orientations diagram from top) conventionally plunged β-gal sample, bottom) β-gal particles from SPOT-RASTR method

We next compared the SPOT-RASTR reconstruction to 3D reconstruction from a conventionally prepared β-gal sample. β-gal was prepared using Vitrobot plunge freezing on Ultrafoil gold grid and cryo-EM data were collected. When the data were reconstructed, inspection of the Euler angles revealed a strong preferred orientation (Fig. 4B top). When the Euler angle plot for the conventionally prepared data (Fig. 4B top) was compared against the SPOT-RASTR data (Fig. 4B bottom), it revealed specimen preparation on the membrane tube filled in most of the missing angles in the Euler plot. The Euler space was not uniformly filled, but we propose this happens due to the particles binding to either side of the tubes more often than binding the top and bottom due to ice thickness. However, in the SPOT-RASTR β-gal data there is nearly even angular coverage along the -π/2 elevation (Fig. 4B, bottom) compared to conventional plunge freezing where there are very few views along that axis. This is likely because the particles rotate freely, so Euler space is still filled even if the particles were only bound to the sides of tubules.

There was a difference in resolution between the map obtained from the SPOT-RASTR method and the map from conventional sample preparation for two major reasons. First, the conventional preparation data is collected in higher mag (smaller pixel size) and the total number of particles are slightly higher. Second, the missing views due to the preferred orientation in the conventional preparation are compensated by the D2 symmetry present in β-gal. The missing views would be more deleterious if it was a sample without any symmetry (Tan et al. 2017).

### SPOT-RASTR provides potential for automated 3D reconstruction

Since the particles bound to the nanotubules have a defined diameter and are located close to the surface of the tubule, we hypothesized that they could be picked by 2D classification, completely automatically without requiring templates or other human intervention. We call this *picking-by-classification*. For this approach, after the tubular segments were picked and tubule signal erased, the segments were split into overlapping sub-boxes where the box size was just bigger than the particle. Thus, the individual particles should be enclosed in one of the sub boxes (Fig.5A, B). The sub-boxes were then classified using rounds of 2D classification. In the first round, the sub-boxes that contained only membrane classified together, the sub-boxes containing nothing formed incoherent class averages, and sub-boxes with particles classified into coherent class averages. The averages that contained particles were subjected to a second round of classification which produced high-quality class averages that represented the full range of particle views present in the SPOT-RASTR sample (Fig. 5D). Thus, the picking-by-classification approach was able identify and center the particles present in the sample and distinguish them from tubule density and noise.

**Figure 5.**
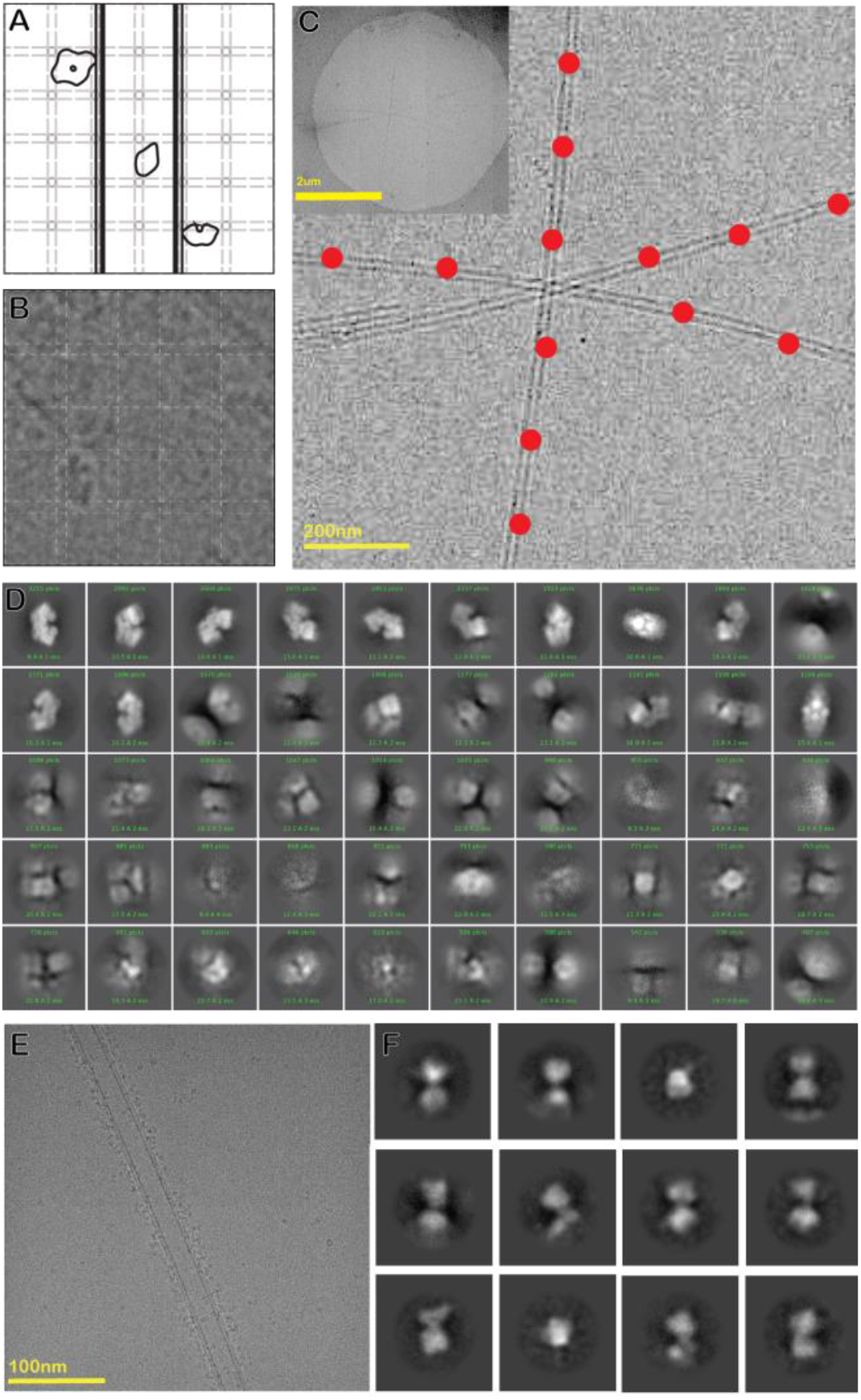
A) Picking by classification scheme. B) An example of picking by classification showing the overlapping picks with dashed lines. C) Picked tubes for TEM acquisitions using a Topaz training model (the hole view is shown in the thumbnail at the top left). D) 2D classes of β-gal obtained from previous section’s picks. E) SiRFP/HP heterodimer decorated on a tube. F) 12 exemplar 2D class averages of SiRFP/HP showing the two globular densities expected for the dimer.

The SPOT-RASTR approach also has the potential for automated data collection without the need for manual selection of low magnification targets. Individual nanotubules can be resolved at the “hole” magnification used for automated data collection (2850 mag, 61.05 Å pixel size). With conventional data collection, holes can be identified by cross correlation or a trained machine learning model (Kim et al. 2023). This approach works well if particles are evenly distributed across holes. However That is frequently not the case (Drulyte et al. 2018; Noble, Dandey, et al. 2018), and filamentous samples can be particularly challenging because they are often not distributed evenly across all holes. However, since the nanotubules can be resolved at lower magnification, data collection can be trained to focus on them exclusively. As a proof of concept, we trained the Topaz picker on manually picked particles. Then we tested the model on 55 low magnification (2800 X, pixel size 61.05 Å) images and it returned 1800 tube particles, skipping empty holes where there were no tubes. This revealed we can target only the tube particles in the hole. Although we have not yet implemented this approach in a data collection software package, our results demonstrate that the tubules can be automatically targeted, and the rest of the grid excluded (Fig. 5C). We estimate that using this approach with ^∼^1 Å/pixel we could get ^∼^150 nonoverlapped tube exposures (high magnification acquisitions) per hour, which will result in ^∼^1700 tube particles picks using filament tracers and then in the case of β-gal, would yield ^∼^1070 particles on average per hour.

### SPOT-RASTR on smaller protein samples

Next, we tested the SPOT-RASTR method with a more challenging sample. The minimal dimer of the sulfite reductase flavoprotein and hemoprotein (SiRFP/HP) is a 110 kDa complex that has a strong tendency to disassemble into its ^∼^50 kDa monomers when plunge-frozen using traditional blotting (Fig. S4). Using the nanotubes, we observed that the Ni affinity tags adsorb most of the free particles to their surface, and we can see only a few free particles outside the tube areas (Fig. 5E). After performing the SPOT-RASTR processing steps for this dataset we obtained 180,000 tube particles out of 5,000 motion corrected micrographs. After subtraction and particle picking, we ended up with 20 2D classes that show defined features (Fig. 5F). The averages have two connected globular densities that are consistent with the 50 kDa monomers and show views that are consistent with a full range of angles for the particle. We did not push these data to 3D reconstruction, because the ice appears too thick for high-resolution reconstruction. Nonetheless, these results show that SPOT-RASTR protected the complex from dissociation into monomers and promotes a full range of views.

## Discussion

Here we have shown that SPOT-RASTR offers several key advantages in addressing the air-water interface issues commonly encountered in specimen preparation for cryo-EM such as partial denaturation, aggregation, and preferred orientation. Additionally, the process has advantages in terms of consistency, reproducibility, and optimization compared to other alternative preservation methods, making it possible to assess its compatibility for a given specific sample rapidly. With SPOT-RASTR we demonstrate a useful and accessible tool for cryo-EM sample preparation that ensures the quality of the particle after freezing. Our results demonstrate that RASTR has the potential to enable cryo-EM specimen preparation that overcomes challenges faced by other methods.

The SPOT-RASTR sample preparation method offers a promising solution for proteins that have a propensity to bind to the air-water interface, resulting in partial or complete denaturation. By utilizing lipid nanotubes, this method effectively addresses the issue by preventing protein binding to the interface. The protein’s interaction with the tubes immobilizes it, preventing its contact with the air-water interface and preserving its structural integrity throughout the blotting and freezing process. Tomograms clearly show the β-gal particles either are bound to the tubes and embedded in bulk ice, or when they are free in solution, they are attached to the air-water interface. This demonstrates the protein’s tendency to bind air/water interface and rationalizes the preferred orientation observation in the conventionally plunge-frozen dataset. Furthermore, comparing the quality of the final particles and resulting map with the conventional plunging method used for the β-gal, it is observed that SPOT-RASTR maintains comparable quality. Thus, the inclusion of additional steps during both plunge-freezing and processing with SPOT-RASTR does not introduce artifacts or otherwise degrade specimen quality.

The protective quality of the SPOT-RASTR method is additionally demonstrated with the SiRFP/HP complex. When prepared using conventional plunge-freezing. SiRFP/HP breaks down into the individual subunits. When prepared with SPOT-RASTR, the sample remains intact, and the dimer interface can be clearly observed in 2D class averages (Fig. 5F). Despite this improvement, the sample still needs more optimization to be reconstructed to high resolution. In this case, we did not collect a sufficiently large dataset, and the ice thickness was likely too thick for high-resolution. We expect that the latter will be the biggest challenge for other samples with SPOT-RASTR. If the tubules stack up on one another (Fig. 2) then that can cause the ice to be too thick ice for high-resolution alignment. In such cases, care must be taken to optimize the cryogenic sample preparation such that it balances tubule density (i.e. tubule concentration) with the resulting ice thickness. Fewer tubules result in thinner ice, but then must be balanced by collecting larger datasets.

The SPOT-RASTR method improves the angular distribution of the particles compared to conventional plunge-freezing for His-β-gal. This enhancement in angular distribution addresses preferred orientation issues arising from interaction with the air-water interface. Compared to the conventional dataset, SPOT-RASTR results in much more even Euler angle coverage, and a nearly complete sampling of Euler space. Tomograms confirm this observation. While there is a random distribution of β-gal particles on the surface of the tubes due to the different His-tag binding from each of the monomers in different angles, normal plunging causes the particles to adsorb into the air-water interface by very limited orientations, which may not be an issue for β-gal reconstruction but have caused a data processing barrier for many datasets. Notably, some of the alternative preservation methods such graphene/graphene-oxide grids or streptavidin monolayer grids can cause the similar preferred orientation. This is because, in the best case, they provide a binding surface and for many reasons it can capture the sample in one or small number of orientations. We have shown that the tubules provide a way around that scenario.

The SPOT-RASTR method exhibits a high level of reproducibility, yielding consistent outcomes in multiple experiments. This held true for both the model protein β-gal as well as a smaller more challenging sample SiRFP/HP. Specimens were successfully captured using the SPOT-RASTR preparation method using different grid types, different blotting conditions, and different concentrations, highlighting the versatility of the method. These parameters can be tuned for each sample to optimize the tubule density, maximize the number of particles decorating the tubes, and achieve preferred ice thickness. SPOT-RASTR feasibility can be assessed at initial stages of preparation through negative stain imaging, where the particles’ decoration is readily distinguishable and verifiable. This is a unique advantage when compared with other alternative preservation methods which may need days or weeks of trial and error to verify their feasibility for the certain sample of interest. Other methods like graphene or affinity grids are comparatively more time consuming and expensive to optimize because the lack of such obvious effects that can be visually obtained and assessed during a simple screening.

Another advantage of the tubules is that they act as miniature affinity column for capture and concentration of His-tagged particles. As a result, the sample concentration that results in ideal tubule binding and decoration is lower than the concentration required for conventional plunge freezing or blot-free vitrification. For example, the β-gal concentration for binding to the tubes after their application to the grid was 0.4 mg/mL, which is lower than 1 mg/mL that was the concentration used for normal plunge freezing. The significance of this is that it allows investigators to work with lower protein sample concentration which is both less expensive and lowers the risk of sample becoming aggregated or denatured during the harsh concentration step. However, it is important to note that different samples may require individual testing and optimization to determine the optimal concentration and binding conditions specific to each case.

We have shown that the SPOT-RASTR approach has the potential to automate the target selection process during data collection, and the particle picking step during data processing. When combined with other cryo-EM tools, these two innovations complete the pipeline for completely automated structure determination. There are numerous software packages for automated data collection including Leginon (Suloway et al. 2005), EPU, and SerialEM (Mastronarde 2003). With each of them, there is typically a laborious step where an experienced user selects targets for data collection or tunes an algorithm for automatically selecting high-magnification targets. However, one does not know the quality of the sample in those holes until after the data is collected. We have shown that individual nanotubules can be automatically targeted at low magnification. Since we know that the specimens of interest will be bound to the nanotubules, by automatically targeting tubules for high-resolution data collection, the SPOT-RASTR approach enables data collection on areas of the grid that have high-quality particles. Once the data are collected, the next step is CTF estimation, and there are many tools available for that (Punjani et al. 2017; Rohou and Grigorieff 2015; Zhang 2016). Typically, the particle picking step requires some human interaction, but our new picking-by-classification technique allows completely automated particle picking and centering. The remaining steps including initial model creation, refinement, and model building each have automated tools available to accomplish them. Taken together, all of these comprise all the steps for automated structure determination. Thus, our new tools demonstrated here provide the potential for fully automated structure determination.

In conclusion, the SPOT-RASTR sample preparation method demonstrates excellent reproducibility, efficient particle binding, high-quality final particles, improved angular distribution compared to normal plunging, and the potential for automation. These advantages collectively contribute to the overall success and reliability of cryo-EM data obtained using this technique, establishing it as a valuable approach for the structural analysis of biomolecules.

## Materials and Methods

### Protein expression and purification

The C-terminus His-tagged β-Galactosidase sequence from *E. coli* was inserted into the pET28 plasmid and transformed into the chemically competent strain BL21(DE3) for protein expression. The quality of the transformation was tested with Western blotting using anti-His antibodies. A single ^∼^130 kDa β-gal monomer band was observed in SDS-PAGE and the blot membrane. The transformed cells were optimized for expression under different parameters like temperature and expression time. Then the cells were transferred into 6L of LB media for large-scale expression. Subsequently, the β-gal protein was purified into 10 mg/mL aliquots using Ni-NTA affinity column and Superose 6 size exclusion chromatography in the 50 mM Tris pH 6.8 and 100 mM NaCl buffer (the elution buffer for the Ni-NTA column contained an additional 500 mM imidazole to elute the β-gal).

SiRFP/HP expression and purification was performed based on a previously published protocol (Murray et al. 2021).

### Nanotube preparation

Lipid nanotubes were made in a 9:1 molar ratio of the GalCer (C24:1 Galactosyl(ß) Ceramide, note the β stereoisomer need to be used, and not the α) and DGS-NTA (18:1 DGS-NTA(Ni)) respectively (Fig. 1). The GalCer lipids were dissolved in 1:1 chloroform:methanol to make 2.5 mg/mL stock solution and the DGS-NTA lipids were dissolved in chloroform to make 25 mg/mL stock solution. The calculation was done considering the 200 μL final solution volume of lipids in buffer, 1 mg/mL total lipid concentration, 9:1 molar ratio of GalCer to DGS-NTA lipids. We first mixed the calculated stock volume of each into a new 2 mL glass container under the hood. Then an Argon gas probe was used to evaporate the solvents in the mixture with a low flow in the glass container, followed by overnight vacuum drying to make sure no solvent is left inside the lipid mixture. The dried lipids were then reconstituted with 200 μL of buffer (50mM Tris and 100mM NaCl). Multiple pipetting cycles were performed to mix the lipids as much as possible, followed by 5 minutes of mild vortexing to obtain the final concentration of 1 mg/mL reconstituted tubes.

### Plunge freezing

For β-gal on tubes, naked membrane tubes were applied on Quantifoil grids coated with gold (2/2 hole sizes, 300 mesh size) while the grid was hung on the Vitrobot Mark IV tweezer and side blotted. Then this step was repeated. Afterward, β-gal at a concentration of 0.15 mg/mL was applied to the grid, and the grid was plunge-frozen into liquid ethane with blot force 1 and blot time of 1s. Then, the grids were clipped to put in the Titan Krios.

For SiRFP/HP on tubes, the tubes were applied to the grid and side blotted in the same way as above. Then the SiRFP/HP at 0.2 mg/mL was applied to the grid and plunged using the same configuration.

For conventional β-gal, the sample was applied to the same type of the grid at 0.25 mg/mL concentration and plunged into liquid ethane with exact same configurations.

### Data collection

Data collection was performed using a 300 kV Titan Krios electron microscope equipped with either Gatan K3 camera or DE-Apollo (shown in Table.1). The Appion/Leginon (Lander et al. 2009; Suloway et al. 2005) data collection software was used to acquire movies. The high magnification acquisitions for both β-gal and SiRFP/HP were targeted manually, and the high-magnification data were collected automatically thereafter.

### Tube alignment and subtraction

The collected frames were subjected to motion correction using Motioncor2 (S. Q. Zheng et al. 2017), and the contrast transfer function (CTF) was estimated using CTFFIND4 (Rohou and Grigorieff 2015).

For the tubes datasets (either β-gal or SiRFP/HP) tube particles were picked using cryoSPARC (Punjani et al. 2017) and filtered by 2D classification for multiple rounds to remove the junk particles. Good particles were then extracted for further processing (numbers are shown at Table.2). RASTR was applied on the extracted particle stack for both SiRFP/HP-tubes and β-gal-tubes datasets. Briefly, RASTR used autocorrelation-based algorithm to find the in-plane rotation angles (psi) of tubular axis in particle images. Specially designed peak finding strategy determined the edges positions of tubes and used them to calculate the minimum shifts needed to place tube at center. Random 0-360 phi angles (around z axis) and 90 or 270 theta angles (around y axis) were assigned to particles. A simple cryo-EM 3D reconstruction using these angles and shifts yielded the azimuthal average map. The map was further smoothed by averaging along z axis (Fig. 3 A, B). RASTR aligned tube particles were then imported into cisTEM (Grant, Rohou, and Grigorieff 2018) and undergone a final global refinement without theta angles. The treated stack were then exported from cisTEM to generate a 3D volume reconstruction using Relion (Zivanov et al. 2018), and this volume was subtracted from the particle stack to remove tube signals in Relion (Fig. 3C, D). The remaining particle stack was treated as micrographs, allowing for individual picking of β-gal particles.

### β-gal and SiRFP/HP particle picking

Two different strategies were employed for particle picking. First, 2D templates created by projecting volumes were used for cross-correlation based particle picking algorithm (in-house script). This resulted in the selection of 145,000 clean particles which after multiple rounds of 2D classification, were selected for the 3D refinement described in the next section. Second, a novel picking by classification method was used, wherein each micrograph was divided into overlapping pieces (Fig. 5A, B). In the β-gal case, 4x4 divisions were used which yielded 16 particles out of the large particle and these pieces were extracted as individual particle boxes to use in 2D classification. This second method yielded approximately 40,000 high-quality particles after multiple rounds of 2D classification.

### 3D reconstruction

An initial model was generated using all the particles in cryoSPARC and subsequently refined with nonuniform-refinement (Punjani, Zhang, and Fleet 2020) with 145,000 final particles result from the template picking approach after multiple rounds of 2D classification to filter the best-aligned particles, resulting in a final resolution of 3.58 Å with D2 symmetry applied.

### β-gal (no tube) particle picking and 3D reconstruction

The β-gal particles were picked using blob picker in cryoSPARC and filtered out with multiple rounds of 2D classification. Then an initial model was built and then refined with nonuniform-refinement in cryoSPARC with 145,00 particles resulting a 2.03 Å resolution map.

### Model refinement

The obtained map was imported in PHENIX package. 3j7h PDB model then used to fit and refine into the 3.58 Å map of β-gal (from tubes data) using the real-space refinement function with minimization global, local grid search and ADP options on.

## Acknowledgements

Funding for this research was provided by National Institutes of Health grants R01 GM148734 and R01 GM148734 to SMS. The SiRFP/HP work was supported by National Science Foundation grants MCB1856502 to MES. Instruments on which the data were collected were supported by NIH grants U24 GM116788, R35 GM139616 and NSF grant 2017869 . Maps and models were deposited to the Protein Data Bank and EM Data Resource: TBD,TBD – conventional β-gal, TBD,TBD – SPOT-RASTR β-gal. Stacks from the tubule datasets were deposited in the EMPIAR database under the accession numbers TBD, TBD.

## Supplemental Information

**Figure S1.**
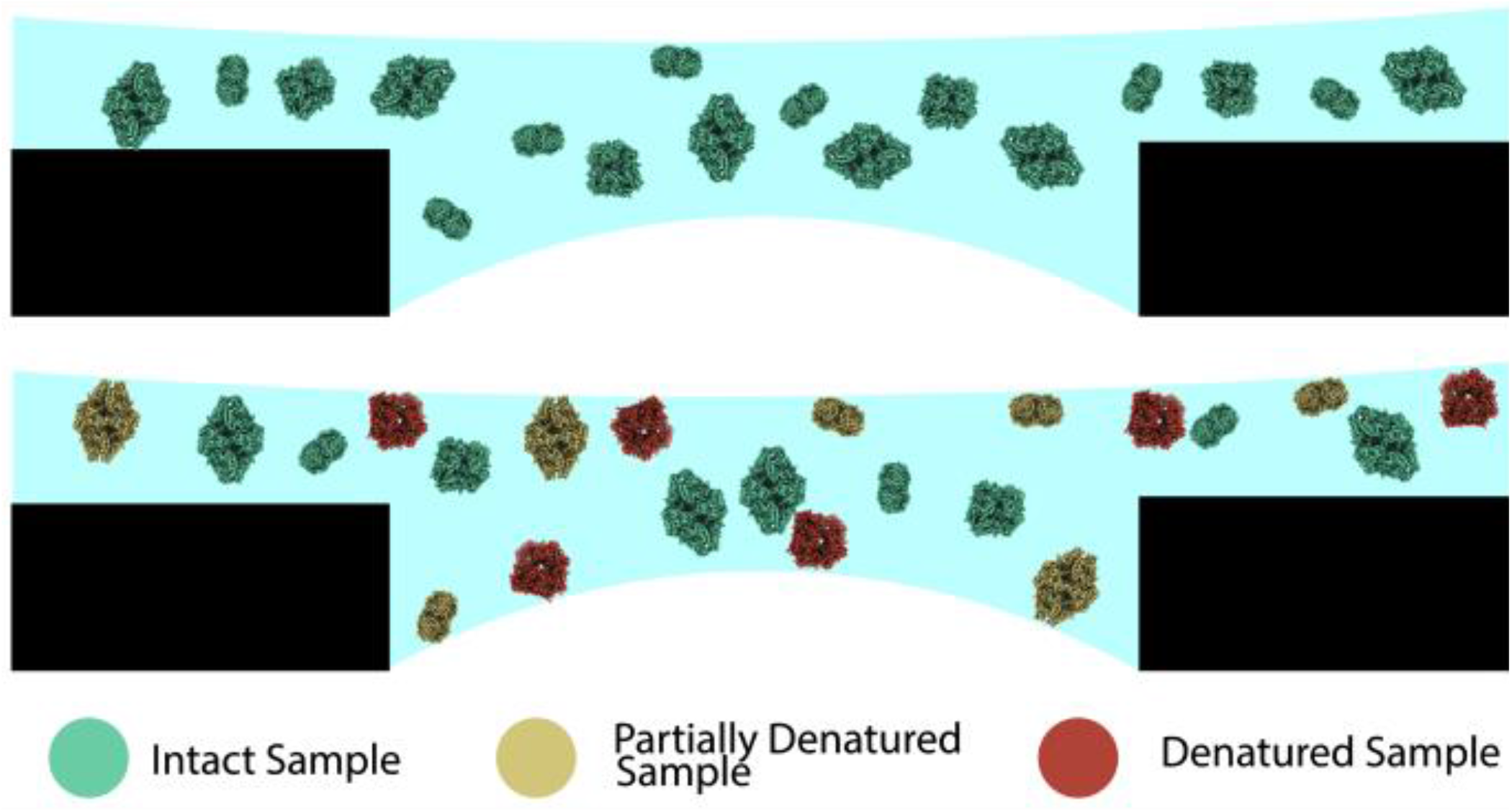
Top) Expected well dispersed β-gal particles in vitreous ice. Bottom: β-gal dispersion in vitreous ice considering the air-water interface absorption of the particles. Bottom) β-gal dispersion in vitreous ice

Video1. Selected tomogram showing two tubes with decorated β-gal.

**Figure S2.**
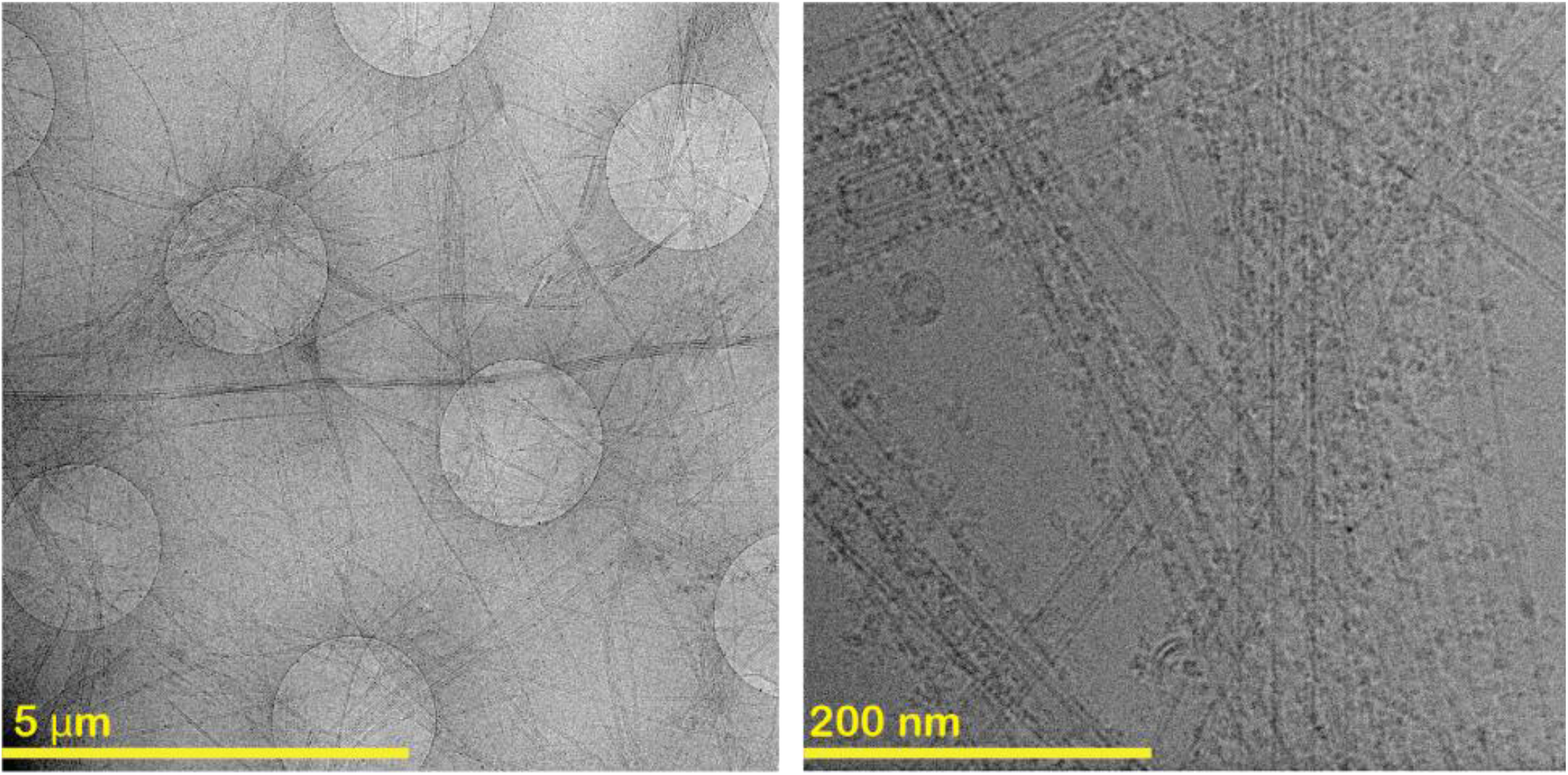
Left) Hole view of the decorated tubes with β-gal showing the huge aggregates due to the premixing. Right: The exposure from one of the previous holes showing the tube aggregate caused by the β-gal multi his-tag

**Figure S3.**
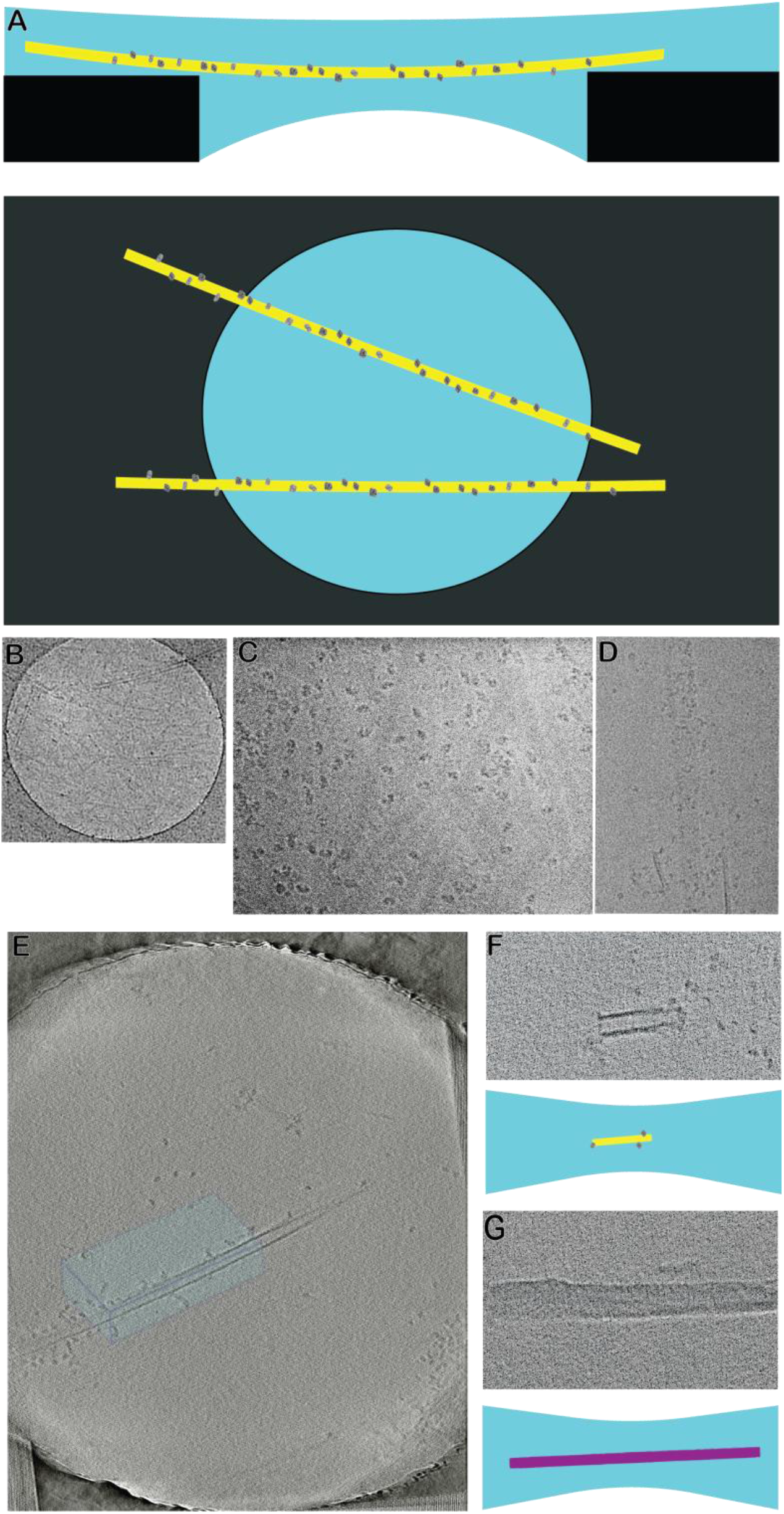
A) Schematic formation of the decorated tube on the cryo-EM grid hole. B) A crowded ghost tubes region hole view. C) Crowded ghost tubes exposure example decorated with β-gal. D) Crowded ghost tubes exposure example decorated with SiRFP/HP. E) A tomogram slice of the decorated tube which its schematic has shown in the Fig.2. F) Short tube decorated with β-gal. G) A dark tube with no decoration.

**Figure S4.**
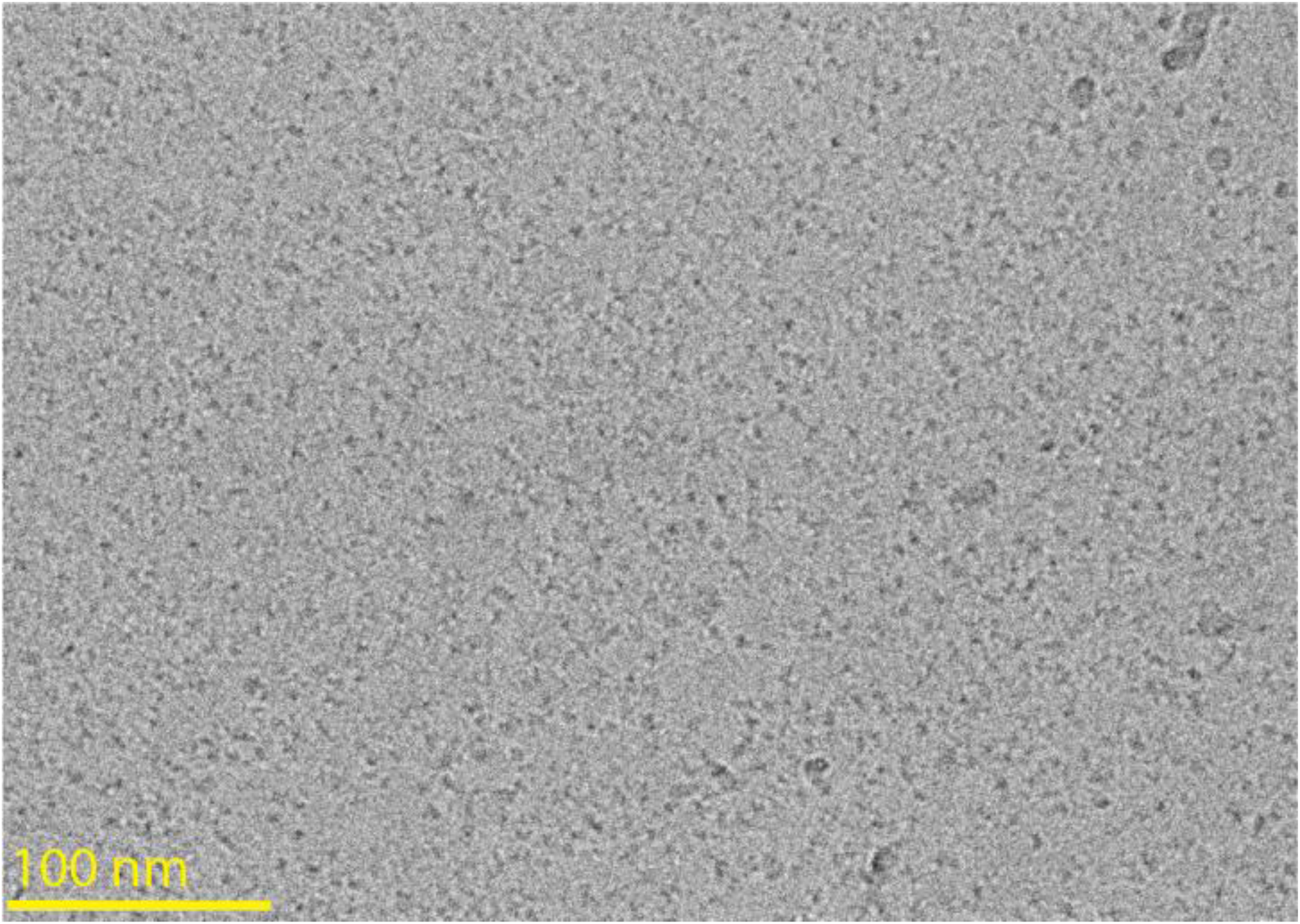
Disassembled SiRFP/HP monomers falling apart, not showing any dimer formation.

