## Supplementary figures and images for "SPOT-RASTR - a cryo-EM specimen preparation technique that overcomes problems with preferred orientation and the air/water interface"

### Video 1

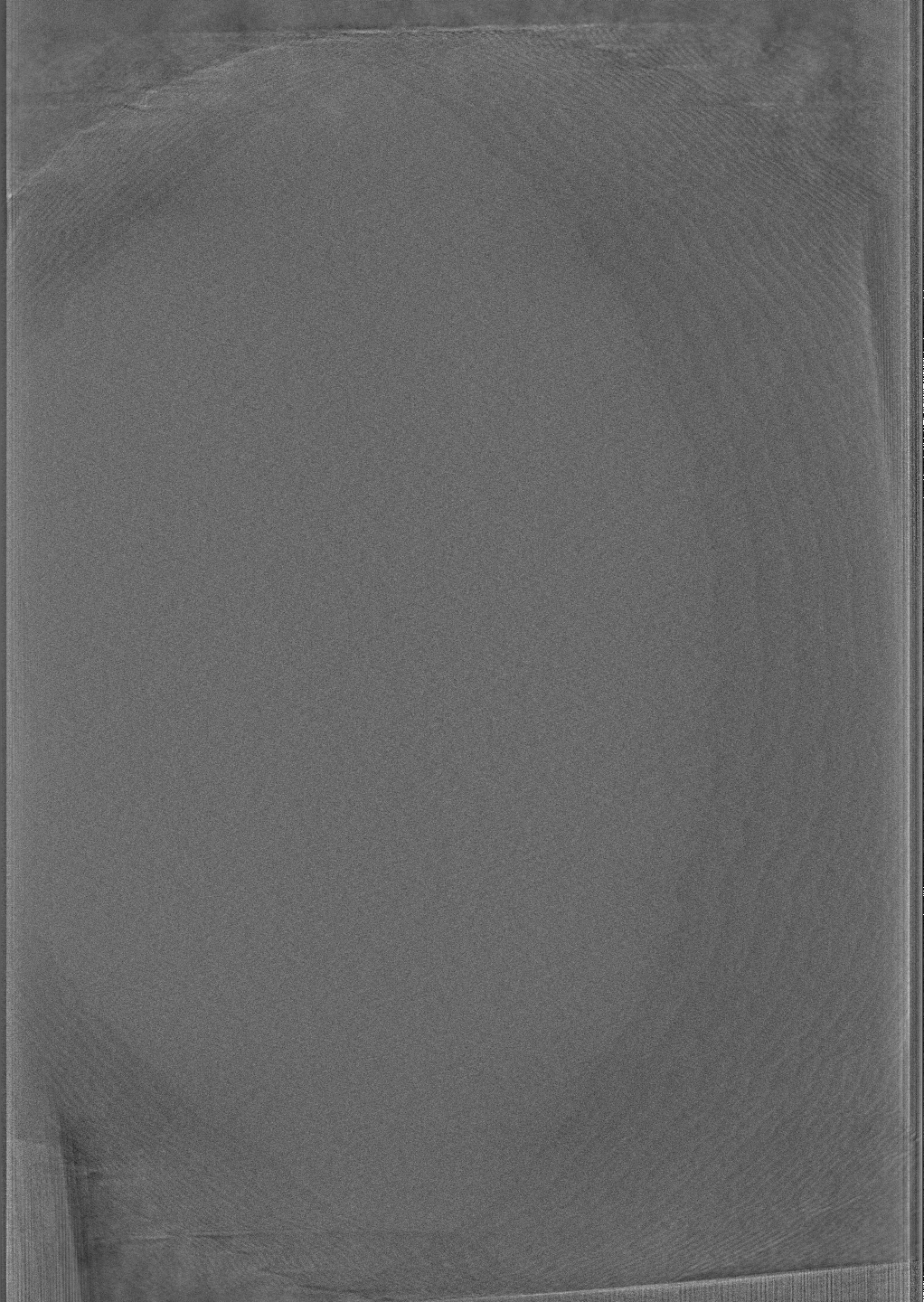
